# Huntingtin fibrils with different toxicity, structure, and seeding potential can be reversibly interconverted

**DOI:** 10.1101/703769

**Authors:** J. Mario Isas, Nitin K. Pandey, Kazuki Teranishi, Alan K. Okada, Anise Applebaum, Franziska Meier, Ralf Langen, Ansgar B. Siemer

## Abstract

The first exon of the huntingtin protein (HTTex1) important in Huntington’s Disease (HD) can form cross-β fibrils of varying toxicity. We find that the difference between these fibrils is the degree of entanglement and dynamics of the C-terminal proline-rich domain (PRD) in a mechanism analogous to polyproline film formation. In contrast to fibril strains found for other cross-β fibrils, these HTTex1 fibril types can be reversibly interconverted. This is because the structure of their polyQ fibril core remains unchanged. Further, we find that more toxic fibrils of low entanglement have higher affinities for protein interactors and are more effective seeds for recombinant HTTex1 and HTTex1 in cells. Together these data show how the structure of a framing sequence at the surface of a fibril can modulate seeding, protein-protein interactions, and thereby toxicity in neurodegenerative disease.

## Introduction

Huntington’s disease (HD) is a debilitating neurodegenerative disease caused by mutant huntingtin with an expanded polyQ region containing more than 35 consecutive Gln residues. Such polyQ expansions render the huntingtin protein and its biologically occurring N-terminal fragments more aggregation prone (Chen et al., 2002) Formation and deposition of aggregated, amyloid-like material from N-terminal fragments of huntingtin is one of the hallmarks of HD. As in other amyloid diseases, a variety of misfolded and aggregated species are formed, but not all of these species contribute equally to toxicity. While some species may be toxic, other species may be less toxic and potentially even protective. This difference is likely to account for findings on the various amyloidogenic proteins, where varying degrees of toxicity have been reported in biological settings, or where different pathways of toxicity appear to be triggered by misfolding. To characterize the roles played by different misfolded species and to foster diagnostic and therapeutic approaches, it is important to understand the structural and functional properties of the various misfolded forms of huntingtin.

Huntingtin exon-1 (HTTex1) is one of the biologically most important N-terminal huntingtin fragments (DiFiglia et al., 1997). HTTex1 contains the disease causing polyQ domain, which is flanked by the N-terminal N17 domain and a C-terminal, proline-rich domain (PRD). Mutant HTTex1 and fragments of similar size are found in diseased tissue and are naturally generated by proteolysis or aberrant splicing (Sathasivam et al., 2013). The importance of mutant HTTex1 in HD is further underscored by animal model studies which show that expression of mutant HTTex1 is sufficient for causing disease symptoms (Mangiarini et al., 1996). Like many other amyloidogenic proteins, mutant HTTex1 forms polymorphic fibrils. In vitro, the yield of a given polymorph is governed by several factors, including temperature (Nekooki-Machida et al., 2009; Lin et al., 2017). It has previously been reported that HTTex1 fibrils grown at 4°C are more toxic than those grown at 37°C (Nekooki-Machida et al., 2009). Both fibril types appear to be present in vivo, but the affected brain regions have been reported to have a higher proportion of the more toxic fibril types (Nekooki-Machida et al., 2009). We have previously investigated the structural features of the more toxic fibril type by electron paramagnetic resonance (EPR) and solid-state nuclear magnetic resonance (NMR). This work has led us to propose the bottle brush model in which the polyQ region forms the β-sheet core region, while the C-terminal PRD projects outward like the bristles in a bottle brush (Bugg et al., 2012; Isas et al., 2015). Cryo electron microscopy of HTTex1 in cellular inclusions supported this model (Bauerlein et al., 2017). The N17 was found to be ordered and in agreement with α-helical structure determined by solid-state NMR (Sivanandam et al., 2011). Different circular dichroism (CD) and infra-red (IR) spectral features suggested structural differences between the fibril types of different toxicity (Nekooki-Machida et al., 2009), but in contrast to the structural information available for the fibrils grown at 4°C and room temperature,(Bugg et al., 2012; Hoop et al., 2016; Isas et al., 2017, 2015; Lin et al., 2017; Schneider et al., 2011; Sivanandam et al., 2011), little is known about the less toxic fibrils grown at 37°C. Knowing the structural features that modulate toxicity could be important for structure-based therapeutic strategies. In addition to a structural characterization, a number of other questions remain unanswered. It is not known whether the different fibril types represent distinctively different strains as seen ubiquitously for other amyloid proteins. Moreover, the seeding behavior of the different fibril types has not been investigated. Seeding plays an important role in misfolding and aggregate formation and it is likely that several types of misfolded proteins can spread within the brain using a seeding mechanism. Thus, in principle different seeding behaviors could have significant impact on aggregate toxicity. In the present study, we combined EPR, solid-state NMR and other biophysical methods to determine the structural differences between fibrils grown at 37°C and 4°C and to test whether they have different seeding abilities in the test tube and in cells.

## Results

It has previously been reported that HTTex1 fibrils at two different temperatures, 4 or 37°C, results in the formation of toxic or non-toxic fibrils, respectively (Nekooki-Machida et al., 2009). In order to characterize the structural differences between those fibril types, we used the established conditions to generate the previously described toxic and non-toxic fibrils from HTTex1(Q46). For simplicity, we refer to these fibrils as T and N-fibrils, respectively. In agreement with the original study, we found that the two different fibril types exhibited distinctively different CD-spectra (Fig. 1a). While T-fibrils typically had a minimum around 213 nm, that of N-fibrils was significantly shifted over to higher wavelengths (218-225 nm). These data are indicative of significant structural differences between the different fibril types. An additional, difference between the two fibril types was their optical appearance. While the T-fibrils were translucent, the N-fibrils were turbid, indicating that these fibrils were larger with a higher propensity to scatter light (Fig. 1b). Consistent with the optical appearance, EM images showed that T-fibrils were mostly non-bundled with typical diameters of 12±2 nm (Fig. 1c) while the N-fibrils were much thicker (220 nm to 10 μm diameter) with a more bundled appearance (Fig. 1d).

**Figure 1:**
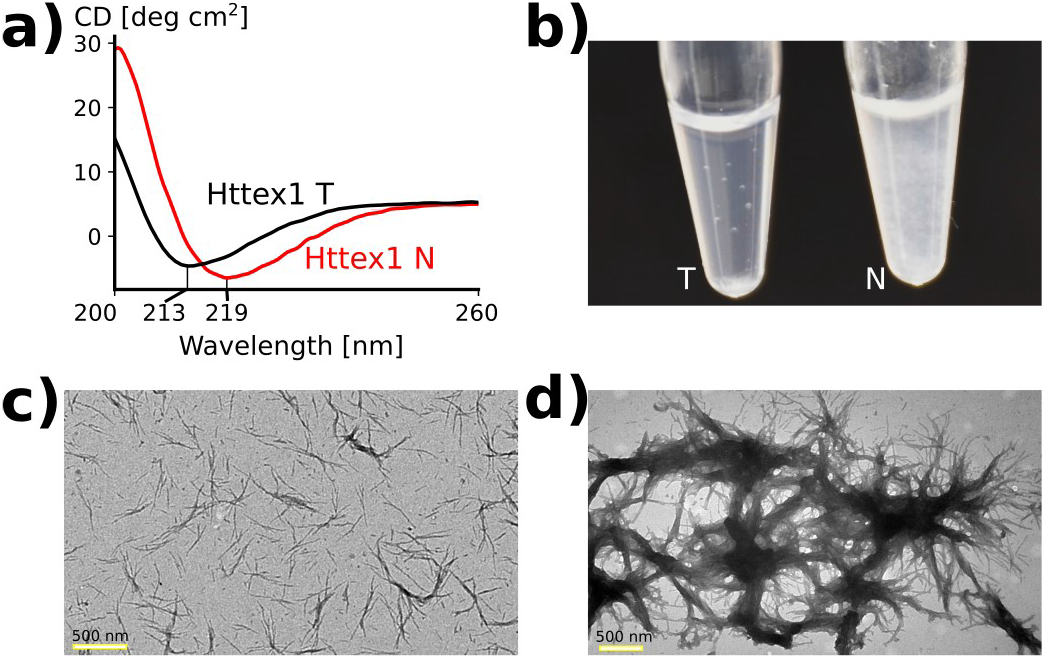
HTTex1 T and N-fibril types are have distinct CD minima, visible appearance, and EM signatures. a) CD spectra of T and N-fibrils are plotted in black and red, respectively. There is a distinct shift in the minimum of the spectrum as reported previously (Nekooki-Machida et al., 2009). b) Visual appearance of both fibrils in suspension. The T-fibrils appear translucent whereas the N-fibrils have a cloudy appearance. c) The EM image of T-fibrils shows little bundeling. d) EM image of N-fibrils shows predominantly bundled fibrils.

We next used a combination of continuous wave (CW) and pulsed EPR as well as solid-state NMR to determine in which regions the structural changes might lie. For the CW EPR experiments, several spin labeled derivatives were generated harboring singly labeled sites within the N17, polyQ or PRD (Fig. 2a). T or N-type fibrils were then grown from these derivatives and the corresponding EPR spectra were recorded. To ensure that proper fibril types were generated, each preparation was tested using CD (Fig. S1). The EPR spectra of T fibrils for sites in the N17 (15R1, 17R1; Fig. 2b, black spectra) and the polyQ region (30R1, 48R1; Fig. 2b) revealed multi-component EPR spectra that were dominated by highly immobilized components. In contrast, the sharp and narrowly spaced spectral lines for position 63 (last residue in polyQ) and the sites in the PRD (71R1, 76R1, 86R1) indicated much higher mobility in this region. These results were very similar to those previously obtained for fibrils grown at low temperature under similar, but not identical conditions (Bugg et al., 2012). Together with our earlier studies, they support the bottle brush model where the PRD bristles face away from the central core of the fibril (Isas et al., 2015).

**Figure 2:**
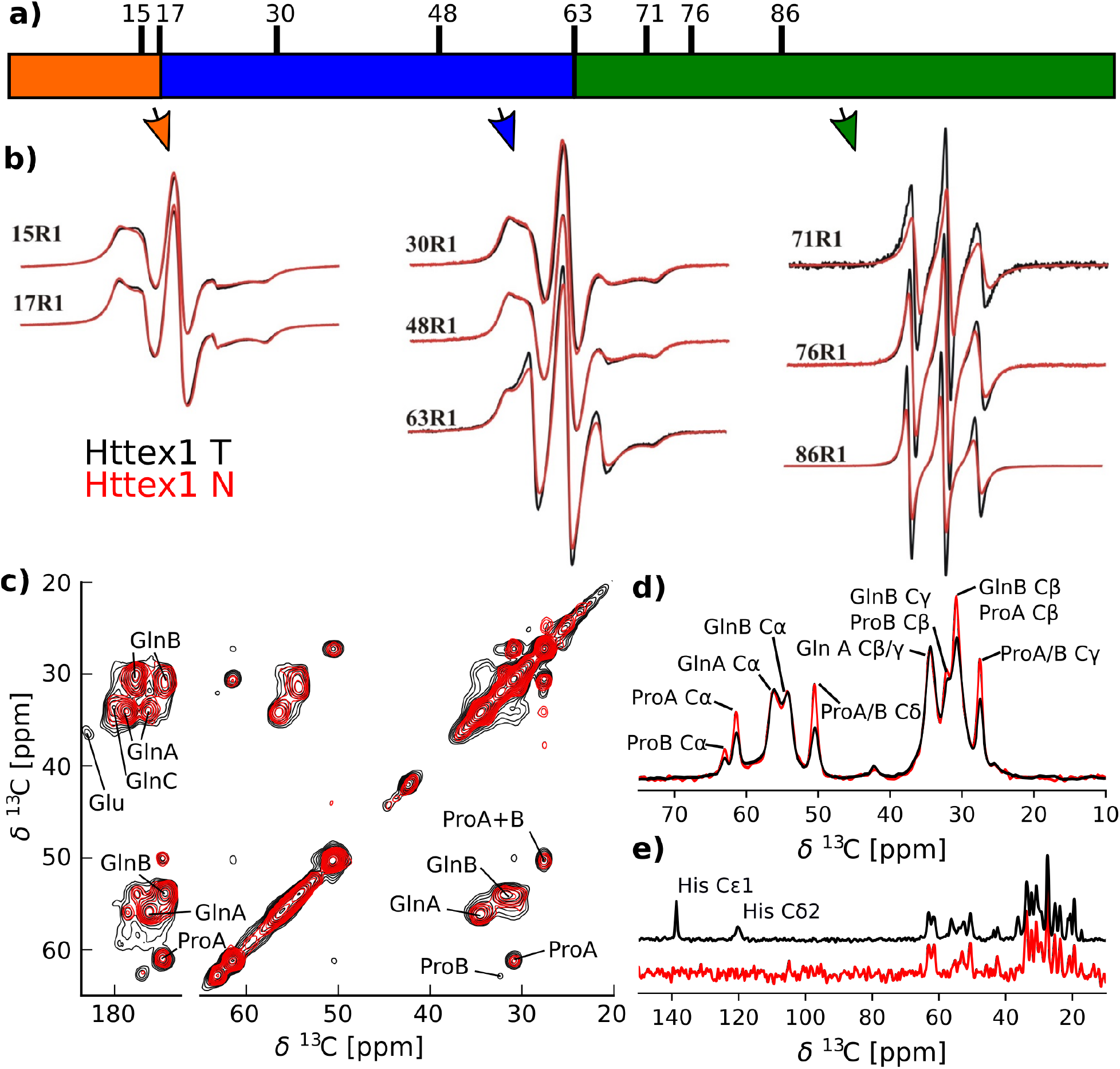
EPR and solid-state NMR spectra indicate a more dynamic PRD in the T-fibrils compared to the N-fibrils. a) Domain structure of HTTex1 highlighting the N17 region in orange, the polyQ domain in blue, the PRD in green, and the C-terminal His-tag in cyan. The residues that were spin labeled for CW measurements are indicated. b) CW EPR spectra of T and N-fibrils are shown in black and red, respectively. The residue number of the spin label is indicated. The spectra of N and T-fibrils overlap very well for all positions within the N17 and polyQ region indicating that they have the same structure and dynamics. However, the spectra of the T-fibrils are narrower for the last Q in the polyQ region (63R1) and all sites within the PRD indicating increased dynamics in this region compared to N-fibrils. c) The 2D DARR ^13^C-^13^C spectra of N and T-fibrils overlap very well, indicating that the static fibril core of both fibrils has a very similar structure. The spectra of N and T-fibrils are shown in red and black, respectively. Both spectra were recorded at 0°C, 25 kHz MAS with 50 ms mixing time. The spectrum of the T-fibrils was previously published as Figure 2 by Isas et al. 2015. d) 1D ^13^C cross polarization (CP) spectra, which are sensitive to more static residues. Spectra of N-fibrils (red) and T-fibrils (black) were normalized to their Gln Cα intensities. The amino acid assignments of the lines are indicated. The higher intensities of the proline lines in the spectra of the N-fibrils indicate that this domain is more static in this fibril type. e) Refocused INEPT spectra, which detect highly dynamic residues, were normalized according to the spectra shown in d). The assignment of the His resonances that were detected for the T but not for the N-fibrils are labeled.

When we compared the EPR spectra from T-fibrils to those from N-fibrils (red spectra in Fig. 2b), we found them to be nearly identical for sites in the N17 and polyQ indicating that the local structure in these regions is highly related in the two different fibril types. Subtle differences, however, were seen at position 63 and these differences became more noticeable for sites in the PRD region. For each of these labeling sites, the spin-normalized EPR spectra of the T-fibrils had higher amplitudes than those of N-fibrils. This was caused by the more pronounced presence of sharp lines in the T-fibril spectra that indicate high mobility. Based upon these data, the T and N-fibrils have related structural features in the N17 and polyQ, but the N-fibrils have more strongly immobilized PRD regions.

To further investigate the differences between the different fibril types, we measured solid-state NMR spectra of T and N-fibrils. To compare the fibril core of these spectra, we used cross polarization (CP) based solid-state NMR spectra that rely on dipolar couplings and that primarily detect the static parts of an amyloid fibril. An overlay of ^13^C-^13^C 2D DARR spectra of T (black) and N-fibrils (red) is shown in Fig. 2c. Both spectra were remarkably similar. Each spectrum contained the two Gln peaks (Gln A and Gln B) that had previously been shown to be the β-sheet structure formed by the polyQ domain (Isas et al., 2015; Schneider et al., 2011; Sivanandam et al., 2011). The difference between Gln A and Gln B have been hypothesized to be the result of alternative side chain conformations (Hoop et al., 2016, p. 2016; Schneider et al., 2011). In addition, cross peaks corresponding to Pro in a polyP II (PPII) conformation (ProA) and weak cross peaks corresponding to Pro in a random coil conformation (Pro B) could be detected in both spectra. All of these peaks superimposed very well in both spectra indicating a highly similar structure. The overall conclusion that N and T-fibril cores have a similar overall structure is also consistent with the 1D CP spectra shown in Fig. 2d, as T and N-fibrils give rise to nearly identical peak positions. However, more detailed analysis of the peak intensities reveals that the Pro peaks are comparatively smaller in the T-fibrils. Inasmuch as the CP spectra are more sensitive to static regions, this indicates that the T-fibrils have a smaller proportion of their Pro residues in a static state.

While CP based spectra focus on the static parts of the fibrils, INEPT based spectra of fully protonated protein at moderate magic angle spinning (MAS) frequencies are selective for highly dynamic domains (Caulkins et al., 2018; Falk and Siemer, 2016; Siemer et al., 2006). The 1D ^13^C refocused INEPT spectra (Fig 2e) show that both T and N-fibrils have considerable dynamic domains. However, a hallmark of the T-fibrils is the presence of His Cε1 and Cδ2 resonances that are not detected in N-fibrils. This difference indicates that the C-terminal His-tag is more dynamic in T-fibrils compared to N-fibrils. Together the 1D CP and 1D INEPT data are consistent with the increased mobility of the PRD observed for T-fibrils by EPR (Fig. 2b).

Having obtained EPR and NMR based evidence that the main differences between the two fibril types reside in the PRD, we next investigated this region structurally by pulsed EPR (DEER)-based distance measurements. Toward this end, we generated four different labeling pairs in the PRD and sparsely incorporated these derivatives (10%) into N or T fibrils grown from excess (90-95%) unlabeled protein. This was done to avoid measurement of intermolecular distances. Two of the labeling pairs, 63-75 and 91-102, flank the two uninterrupted polyP repeats P11 and P10, respectively. The 75-91 pair borders a Pro-rich linker region L17, which connects the two polyP repeats. The 63-102 labeling pair was designed to obtain distance information encompassing the entire PRD (Fig. 3a).

**Figure 3:**
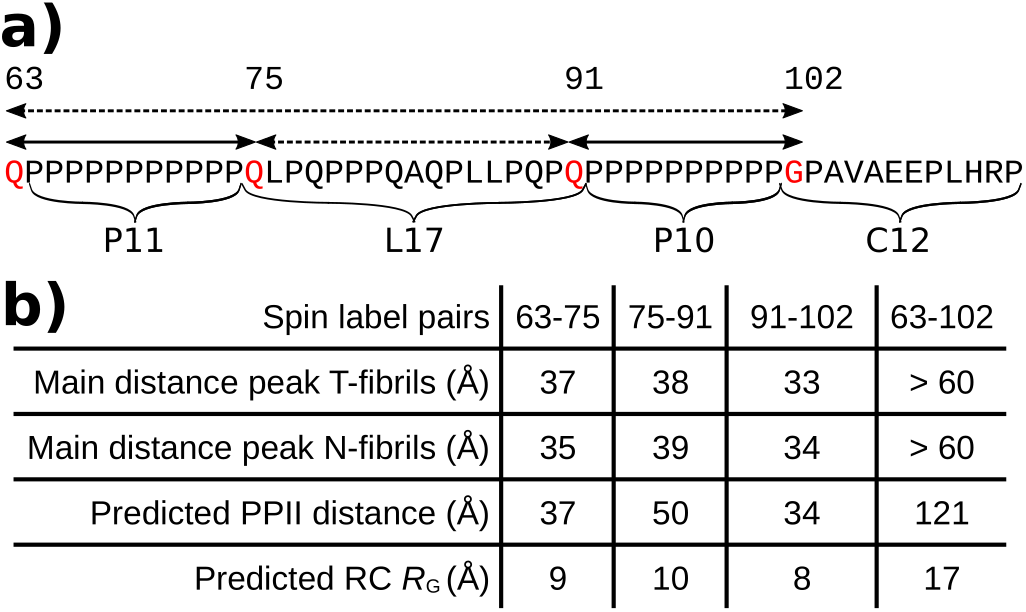
Long-range distance inside the PRD of N and T-fibrils are highly similar. a) Sequence of the PRD indicating the positions of the labels introduced for distance measurements in red and the distance pairs as arrows. The for segments of the PRD including the two polyP regions (P11 and P10) and the linker region (L17) and C-terminals region (C12) are also indicated. b) Table listing all distances measured via EPR DEER experiment for each spin label pair and fibril type. The distance corresponds to the maximum of the main peak derived from the DEER decay curve (DEER data and fits are shown in Fig. S2). In addition, the theoretical distances for a polyP II helix assuming an increase of 3.1 Å per residue and the theoretical radius of gyration (*R*_G_) for a random coil, are given (*R*_G_=*R*_0_ *N^ν^* with *R*_0_=1.927 Å, ν=0.598, and N is the number of residues). Both distances spanning the polyP regions (P11 and P10) correspond nicely to a PPII distance. The distance spanning the L17 region is significantly shorter than a PPII helix and longer than expected for a random coils structure.

We first investigated the distances of the T-fibrils. As expected for regions that contain large amounts of PPII helices (Radhakrishnan et al., 2012), the distance distributions for the 63-75 and 91-102 labeling pairs are broad (Fig. S2). As summarized in Fig. 3b, the peaks of the distance distributions (37 and 33 Å, respectively) correspond remarkably well to the predominant, fully extended form of a PPII helix which increases in length by ~3.1 Å/amino acid (37 and 34 Å, respectively). These distances are not consistent with a random coil structure that would, from theoretical considerations, result in much shorter distances (Kohn et al., 2004). The distance for the 75-91 labeling pair (38Å) was also much longer than expected for random coil structure, but it was also significantly shorter than expected for a perfect PPII helix (50 Å). Together with the solid-state NMR data, which revealed a mixture of PPII helical and random coil peaks for proline residues in the PRD (Caulkins et al., 2018; Isas et al., 2015), this indicates the formation of a less than fully extended PPII helical structure between residues 75-91. Despite this imperfect PPII helical structure between residues 75-91, we found no evidence for a folding back of the PRD onto itself, as the distance for the 63-102 pair was very long, beyond the limit of detection (~60 Å), indicating that residues 75-91 introduce only a slight kink. Interestingly, highly similar distance distributions for the respective labeling pairs were also observed in N-fibrils, indicating that overall conformations of the PRD bristles are similar. Collectively, the solid-state NMR, CW and DEER EPR data indicated that the overall structure in the PRD region was similar in N and T-fibrils with significant amounts of PPII helical structure present in both cases. The central difference between the two fibril types, however, resides in the degree of packing interactions in the PRD bristles. While these bristles freely radiate outward in T-fibrils as shown previously (Bugg et al., 2012; Isas et al., 2015), they are much more tightly entangled in the thicker and more bundled N-fibrils. Considering that the two fibril types are structurally highly similar except for the thickness of the fibrils and the packing of the bristles, it may be possible that N-fibrils are made up from T-fibril building blocks that are held together by interactions between their PRD bristles.

To further test the notion that proline rich regions could mediate protein-protein interactions, we investigated the oligomerization properties of polyP peptides using CD experiments. At low temperatures, the polyP peptides exhibited a characteristic CD signature for a PPII helix with a minimum at 207 nm and a small, but detectable maximum near 230 nm (Fig. 4a, black line). Upon heating, an entirely different spectrum with a minimum near 220 nm appeared (Fig. 4a, blue line). Similar transitions have been reported and interpreted in terms of temperature dependent oligomerization or film formation of the polyP peptides (Tooke et al., 2010). To test for film formation in our polyP preparations, we removed the liquid from the CD cuvette at high temperature and recorded the CD spectra (Fig. 4a, red line). Much of CD spectrum remained after liquid removal indicating that the peptides no longer resided in solution but rather formed a film on the surface of the glass (Tooke et al., 2010). Interestingly, polyP film formation causes a red shift in the minimum of its CD spectrum that is reminiscent of the difference between T and N-fibrils. To investigate these potential similarities further, we generated the difference spectra for the polyP peptides at high and low temperature as well as T and N-fibrils (Fig. 4b). The respective difference spectra have overall similar shapes with maxima near 206 nm and minima near 223 nm. These results are consistent with the notion that the entangeling of PRD bristles could be linked to the generation of N-fibrils.

**Figure 4:**
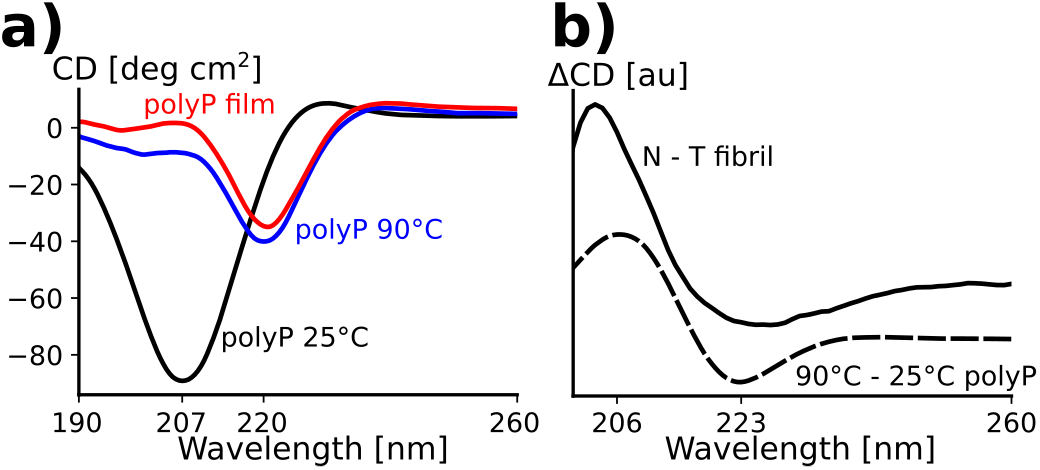
PolyP and HTTex1 difference spectra suggest an increased entanglement of the PRDs in N compared to T-fibrils. a) CD spectra of polyPro at a concentration of 0.1 mg/ml recorded at 25°C (black) and 90°C (blue). The spectrum in red was recorded after the spectrum at 90°C and after removing the liquid from the CD cuvette indicating the formation of a polyP film inside the cuvette. b) Difference of N and T-fibril CD spectra of Fig. 1 (solid line) and polyP CD spectra recorded at 90°C and 25°C (dashed line). The shared maxima at about 206 nm and minima at about 223 nm indicate that the structural differences that are occur during heat-induced polyP film formation are comparable to those between HTTex1 N and T-fibrils.

The overall similarity between T and N-fibrils and the propensity of polyP region to self-aggregate suggested that N-fibrils might arise from T-fibrils via entanglement of PRD bristles. To investigate this hypothesis further, we generated T-fibrils and followed their long-term stability over time. As shown in Fig. 5a, the CD spectra slowly changed over time with the minimum shifting to larger wavelength, eventually approaching the CD spectrum of N-fibrils after 9 days at 4°C. The change in the CD spectra coincided with reduced amounts the sharp (mobile) component in the EPR spectra of 81R1 (Fig. 5b) as expected for a transition to N-fibrils. We also re-measured the 1D ^13^C refocused INEPT solid-state NMR spectrum on the same sample of T-fibrils after 12 months and found that the of His Cε_1_ and Cδ_2_ resonances became undetectable similar to the corresponding spectrum of N-fibrils (Fig. 5c).

**Figure 5:**
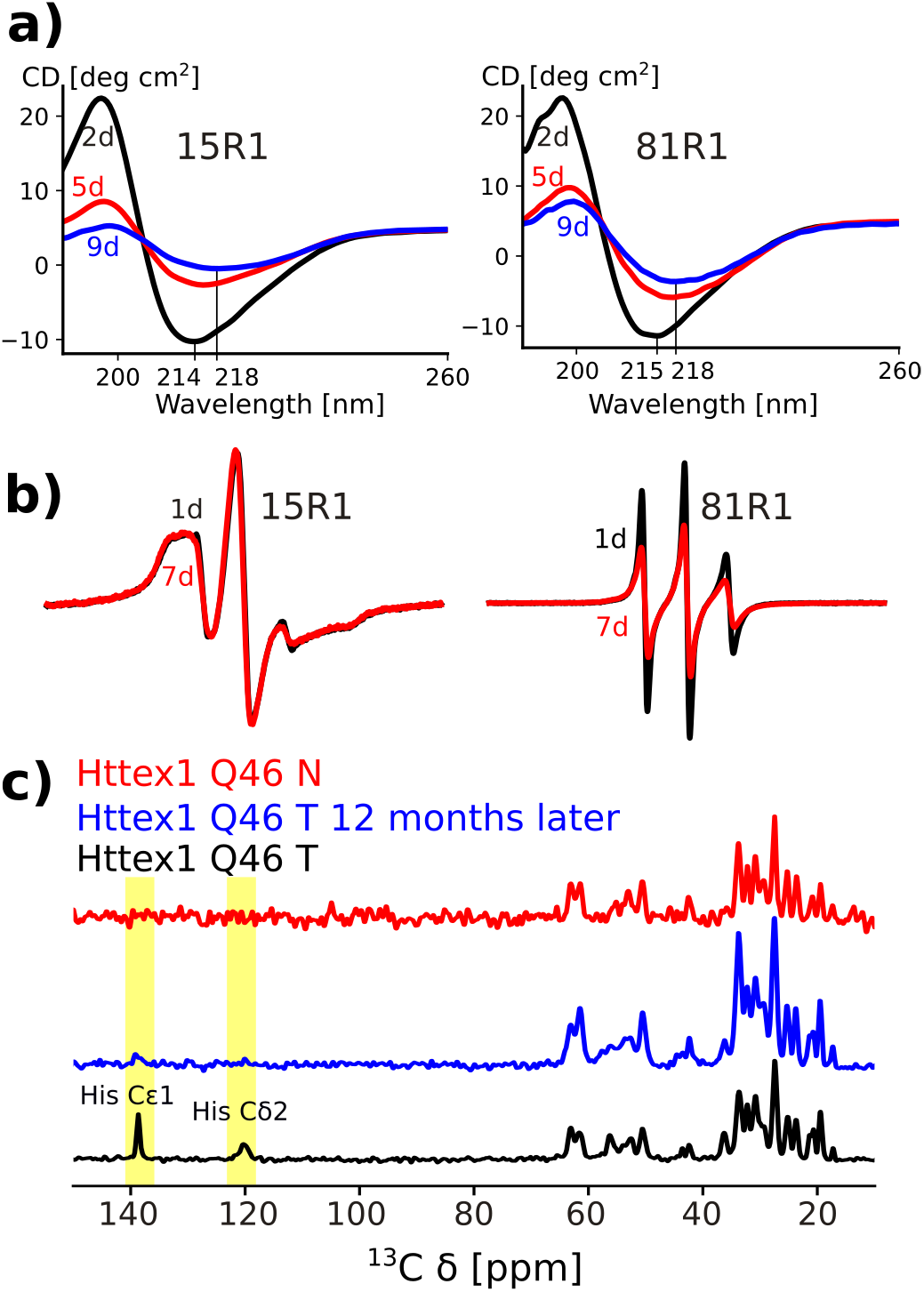
Over time T-fibrils turn into N-fibrils. a) CD spectra of HTTex1(Q46) fibrils that were spin labeled at the N-terminus (15R1) and the PRD region (81R1). Spectra of fibrils incubated at 4°C for 2 days (black), 5 days (red) and 9 days (blue) are shown. The decrease in apparent CD and the blue shift of the minima are compatible with a transition from T to N-fibrils. b) CW EPR spectra of corresponding fibril preparations after 1 day (black) and 7 days of incubaiton at 4°C (red). The spectrum of the N-terminal 15R1 stays unchanged, whereas the spectrum of 81R1 decreases in intensity over time indicating a decrease in mobility. c) Refocused INEPT spectra of HTTex1(Q46) T-fibrils (black), the same fibrils 12 months after fibrillization (blue), and N-fibrils (red). The His resonances, which are detected in freshly prepared T-fibrils, disappear after 12 month and are neither visible in N-fibrils.

Having established that T-fibrils can slowly transition into more entangled, N-like fibrils, we were wondering whether the transition from T to N-fibrils could be reversed. We found that treatment of N-fibrils with 0.5% TFA in H_2_O yielded a strong blue shift in the CD spectra of N-fibrils (Fig. 6a), a transition characteristic for the formation of T-fibrils (Fig. 1a). These data further supported that N-fibrils are formed from T-fibrils through the entanglement of the PRD region, which can be reversed by treatment with organic solvent. Interestingly, we noticed that treatment of N-fibrils with HFIP followed by 0.5% TFA resulted in progressively blue shifted minima in the CD spectra (Fig 6a). This change was accompanied by the formation of increasingly disaggregated fibrils (Fig. 6c). Thus, this treatment disaggregated N-fibrils into fibrillar structures that were shorter and even less bundled than T-fibrils. Inasmuch as we also saw the formation of such short fibrils during early time points of T-fibril formation, we refer to them as protofibrils or P-fibrils. If N-fibrils can be successively converted into T and P-fibrils, one would expect that directly applying the HFIP/TFA treatment to T-fibrils should result in P-fibrils. Indeed, this is the case as shown in Fig. 6d.

**Figure 6:**
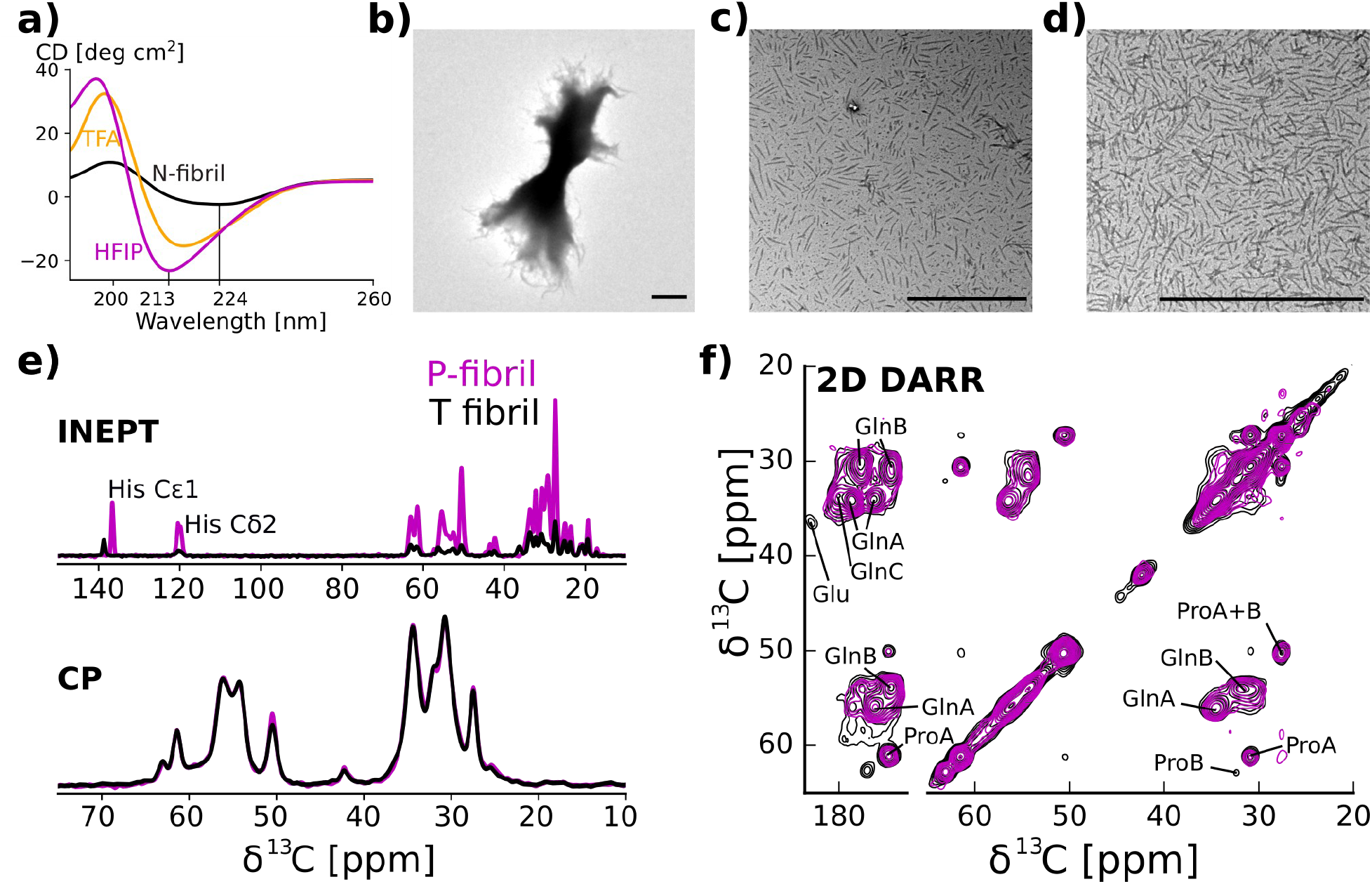
TFA and HFIP can disaggregate T-fibrils to form highly disentangled P-fibrils. a) CD spectra of N-fibrils (black), N-fibrils that were treated with 0.5% TFA in H_2_O (orange), and HFIP (purple). The minima of the spectra treated with 0.5% TFA and HFIP are increasingly blue shifted and show an increase in apparent circular dichroims due to a decrease in scattering. b) EM of highly bundeled N-fibrils before HFIP treatment. c) EM of same fibrils after HFIP treatment showing the presence of highly unbundled proto fibrils. e) EM of N-fibrils that after HFIP/TFA treatment. Scale bars denotes 1 μm. e) 1D ^13^C spectra of HTTex1(Q46) P-fibrils and N-fibrils fibrils are shown in violet and black, respectively. CP spectra were normalized to their Gln C_α_ intensities. The close to perfect overlap of the CP spectra indicates that the static core of these two fibril types is very similar. Refocused INEPT spectra were normalized according to the CD spectra. His side chain carbon resonances are indicated. The much higher relative intensity of the INEPT spectra indicates that the PRD is more dynamic. The shift in the His C_ε1_: line reflects the low pH used for the protofibril preparation. f) overlap of 2D DARR ^13^C-^13^C spectra of HTTex1(Q46) P-fibrils (violet) and T-fibrils (black) indicates that the static fibril core of both fibrils has a very similar structure. Both spectra were recorded at 0°C, 25 kHz MAS with 50 ms mixing time. The spectrum of the 4°C fibrils was previously published as Figure 2 by Isas et al 2015.

In order to structurally characterize the P-fibrils, we used solid-state NMR. Fig. 6e shows the comparison of 1D ^13^C refocused INEPT and CP spectra of HTTex1 T-fibrils and P-fibrils. The CP spectra, which were normalized to the intensity of the Gln Cα peaks, overlap perfectly indicating that the structure of the static fibril core of these two fibril types is very similar. This is confirmed by the overlap of the 2D DARR spectra shown in Fig. 6f. However, the intensities of the refocused INEPT spectra, which were scaled according to the CP spectra, are different. The higher overall signal observed for the P-fibrils, indicates that the C-terminal domain in these fibrils is more dynamic than in the T-fibrils. Except for the shift of the His Cε1 line seen in the refocused INEPT spectrum, no chemical shift change was observed between the N, T, and P-fibrils (see also comparison of INEPT HETCOR spectra in Fig. S3).

Do the fibril types described above show a different degree of interaction to known HTTex1 fibril binding partners? To answer this question, we used dot blots with the MW8 antibody and an anti-polyHisitinde antibody as control. MW8 recognizes an epitope in the PRD and is known to exclusively bind to HTTex1 fibrils (Baldo et al., 2012; Ko et al., 2001). Removing this epitope by replacing the Pro-rich linker in the PRD with prolines, abolishes the binding of MW8 to HTTex1 (Ko et al., 2018). As can be seen from Fig. S4, MW8 binds strongly to P and T-fibrils, less efficient to N-fibrils, and does not bind to the soluble HTTex1 fusion protein or the HTTex1 mutant lacking the MW8 epitope (HTTex1 Pro). Interestingly, dissociating N-fibrils via sonication similar to the production of HTTex1 seeds, increase MW8 binding. Together, these data indicate that less entangled and more dynamic PRDs allow the fibril to engage in more protein-protein interactions, which could explain their higher toxicity.

Seeding has long been considered an important aspect by which the misfolding of amyloidogenic proteins is rapidly propagated. We therefore tested the differential ability of P, T and N-fibrils to act as seeds. We performed seeding reactions using recombinant HTTex1 and used the previously developed time-dependent change in EPR signal of HTTex1 labeled at position 35 as a readout (Pandey et al., 2018). As shown in Fig.7a and b, all fibril types enhanced the loss in signal amplitude, indicating that they all were able to act as seeds. The seeding efficiency was most pronounced in the case of P-fibrils, while that of T-fibrils was weaker and that of N-fibrils was weakest.

**Figure 7:**
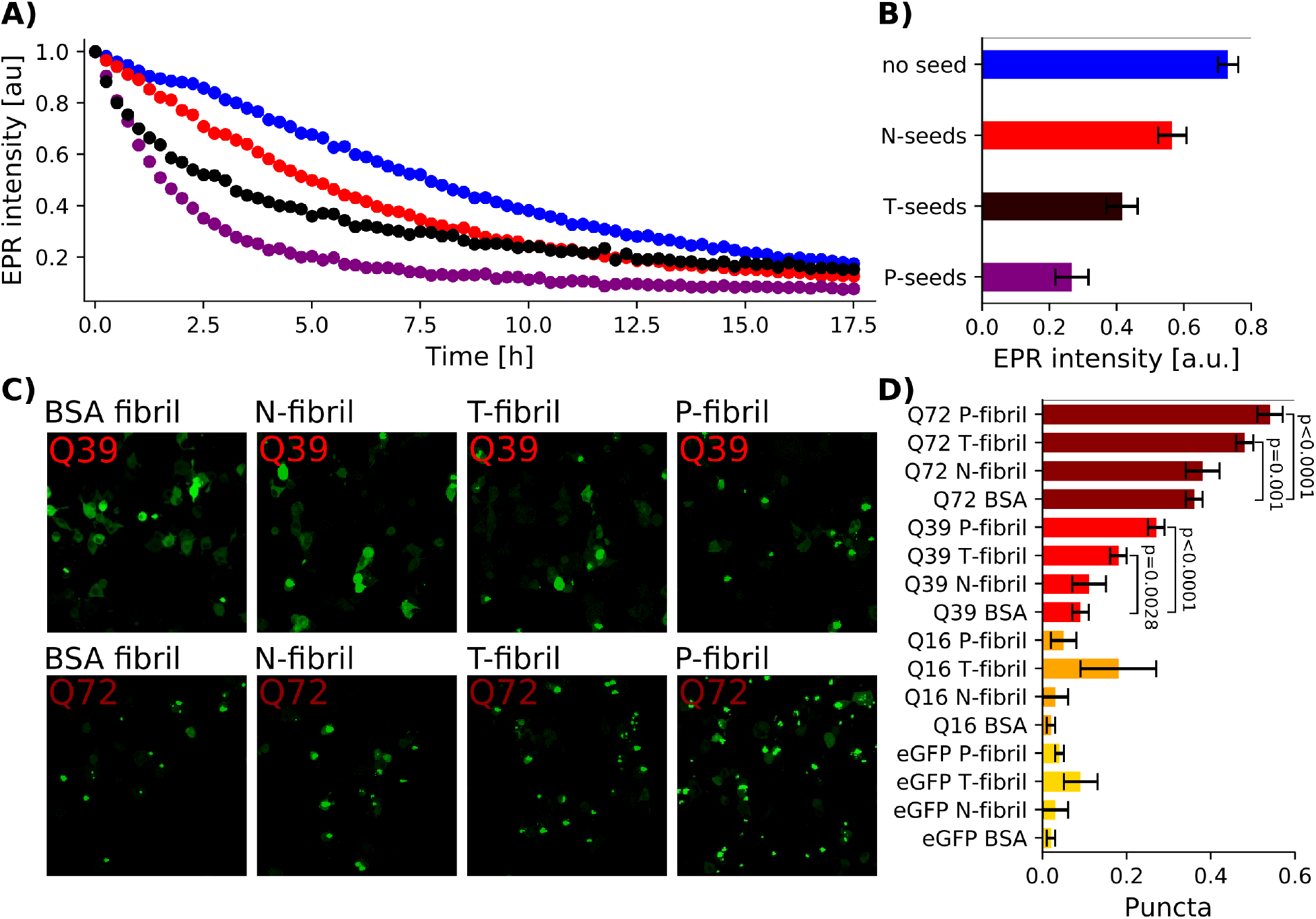
HTTex1 fibrils with disentangled PRDs are more efficient in seeding soluble HTTex1. a) Time dependent CW EPR intensities of unseeded HTTex1(Q46) 35R1 and HTTex1(Q46) 35R1 seeded with N, T, and P-fibrils. b) EPR intensity of HTTex1(Q46) 35R1 4 h after initiating fibril formation averaged over 3 separate experiments. The error bars indicate the standard deviation. The decrease in intensity, which is a reporter of fibril formation, is fastest when seeding with P-fibrils followed by seeding with T, N-fibrils, and unseeded protein. c) Puncta formation in Neuro2a cells transfected with HTTex1 with varying polyQ length, and a C-terminal EGFP tag and EGFP as control. Cells were co-transfected with N, T, and P-fibril seeds and BSA fibril seeds as control. Fluorescence microscopy images were taken 48 after transfection. d) Fraction of transfected cells containing puncta depending on HTTex1 polyQ length and nature of co-transfected fibrils seeds. Experiments were repeated three separate times. Error bars indicate the standard deviation. Significant P-values from a two sided t-test are indicated. P-fibril seeds led to the strongest induction of puncta in cells transfected with HTTex1Q72 and Q39 followed by T-fibrils. The effect of N-fibril and BSA fibril seeds on puncta formation was roughly the same. Few puncta were observed for HTTex1Q16 and EGFP independent of co-transfected fibril seeds.

To confirm these results in the context of cells, we transfected Neuro2a cells with HTTex1-EGFP harboring different Q-lengths (Fodale et al., 2014) and tested how exogenously added fibrils promote aggregate formation of endogenous HTTex1-EGFP within the cells. As shown in Fig.7 c and d, P-fibrils were most potent at seeding as revealed by the strongly enhanced propensity to form puncta for cells expressing HTTex1-EGFP Q72 or Q39. This effect was stronger than for the T-fibrils, which, nonetheless, still exhibited strong seeding ability. In contrast, N-fibrils had a poor seeding ability as puncta formation for these fibrils was comparable to that of the unseeded case. As controls, we also used cells expressing HTTex1-EGFP with a short Q-length (Q16), which are known to have a low fibril forming propensity or EGFP alone. Neither of these cells exhibited any detectable change in puncta from treatment with the different fibrils indicating that seeding effect, which was quantified by puncta formation, is specific to HTTex1 with expanded Q-lengths. In summary, our data show that less entangled PRDs result in fibrils that can seed soluble HTTex1 much more potently. The ability to propagate more efficiently is another important aspect that help explain the higher toxicity of T-fibrils.

## Discussion

Using a combination of CW and pulsed EPR, solid-state NMR, CD and EM, we investigated the structural features of three different fibril types, two of which were previously found to have very different toxicities. Despite the different CD spectra of the fibril types, the overall architecture of their respective core regions was remarkably similar according to CW EPR and solid-state NMR. While the structures of the individual PRD bristles were also very similar, their dynamics varied among the different fibril types. T-fibrils and P-fibrils exhibited the characteristic bottle brush structures where bristles have high mobility and radiate outward. In contrast, the bristles in N-fibrils are less dynamic and EPR, solid-state NMR and CD supports the notion that polymerization of the PRD bristles is involved in the formation of N-fibrils by coalescence of T-fibrils or P-fibrils (Fig.8). This model is furthermore consistent with the greater diameter of N-fibrils and the fact that all fibril types can be reversibly interconverted.

**Figure 8:**
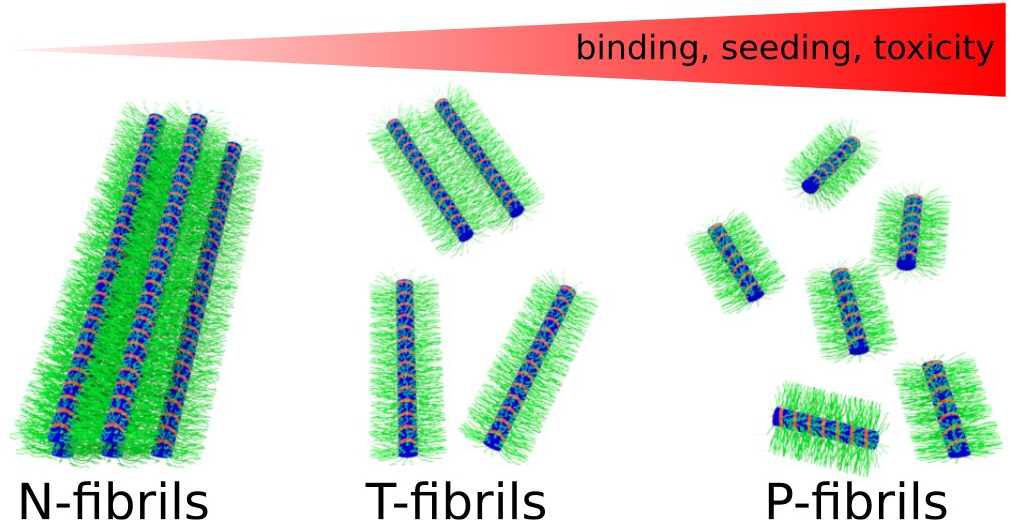
Model of the N, T, and P-fibrils. The PRDs of the N-fibrils (green) are entangled resulting in bundling and consequently less accessible fibril surfaces. The PRDs of the T-fibrils are keeping the individual fibrils more separate. The P-fibrils are shorter and their PRD are even less restricted and more accessible.

The finding that HTTex1 can exist in a variety of different fibril types is a property that is shared with the vast array of amyloidogenic proteins for which fibril polymorphs have been described (Bousset et al., 2013; Gath et al., 2014; Kodali et al., 2010; Lu et al., 2013; Qiang et al., 2017; Safar et al., 1998). What is unusual about the HTTex1 fibril types is that they are different from strains that have been observed in the other amyloidogenic proteins. Such strains typically vary in the basic underlying structures within the core region. In contrast, HTTex1 fibril types are interconvertible, being able to transition from P to T and then to N-fibrils in a process that can be reversed by treatments with organic solvent. This treatment rapidly reverts the CD and EPR spectra of the PRD from N into T-fibrils, without significantly affecting the EPR spectra and solid-state NMR signals of the N17 and polyQ domains. Such a behavior is different from the various strains found in Aβ and α-synuclein where the architecture of the core region is faithfully propagated by seeding and the fibrils need to be completely disaggregated into monomers to transition from one polymorph to another. Thus, the different fibril polymorphs of HTTex1 do not exhibit the typical strain behavior seen in other amyloids.

HTTex1 consists of three separate domains of which the N17 and the polyQ are typically considered to promote aggregation (Jayaraman et al., 2011; Pandey et al., 2018), whereas the PRD is sometimes considered to inhibit aggregation (Crick et al., 2013; Darnell et al., 2007). Here we find that all domains partake in aggregation, but the time scales and structures are vastly different. Prior kinetic experiments demonstrated that the N17 initiates oligomerization on the minutes time scale and subsequently the polyQ regions engages in β-sheet formation on an hour time scale (Pandey et al., 2018). Here we find that the PRD also contributes to self-association as it promotes the coalescing of fibrils. This process is much slower than the N17 or polyQ dependent aggregations, only proceeding on the days time scale. Moreover, each of the domains uses a different secondary structure to promote aggregation. The N17 forms oligomers via helix-helix interactions, the polyQ forms fibrils via β-sheet structures, and the PRD leads to fibril-fibril interaction via the entanglement of PPII containing domains. The latter adhesive interactions are related to those formed by polyP peptides which can form insoluble oligomers and films. All three domains and their interaction are of potential therapeutic interest. While blocking of the misfolding interactions of the N17 and polyQ region could be used to prevent fibril or oligomer formation altogether, it could be beneficial to promote the interactions between PRD region considering that the T-fibrils are thought to be more toxic than the more entangled N-fibrils.

If the structural differences among the different fibril types are directly linked to toxicity, one would also expect that the PRD could be a contributor to HTTex1 toxicity. Intrabodies or proteins such as profilin, which bind to the PRD region have been shown to reduce HTTex1 aggregation and toxicity (Southwell et al., 2009, 2008; Burnett et al., 2008; Shao et al., 2008; Posey et al., 2018). One of the factors that could contribute to the different toxicities of the HTTex1 fibril types is their seeding ability. Our in vitro studies show that the toxic T-fibrils are much more capable of acting as seeds than the non-toxic N-fibrils. This feature is consistent with the finding that the least seeding competent N-fibrils are also least toxic (Nekooki-Machida et al., 2009). Interestingly, the P-fibrils, which have the least degree of entanglement are the most potent seeds. Thus, seeding ability seems inversely related with bundling of the fibrils. Similar results were obtained using Neuro2a cells, where exogenously added fibrils were added to seed endogenously expressed HTTex1. Seeding ability has been considered an important factor in toxicity and cell-to-cell spreading of toxic species in a number of amyloid diseases. Our results therefore indicate that increased entanglement of the PRD protects from seeding and spreading of toxic species.

In addition to modulating seeding activity, the PRD region could further affect toxicity through protein-protein interactions. Huntingtin is a major hub for protein-protein interactions and the PRD plays a central role in many of these interactions because many huntingtin binding proteins contain SH3 domains and other PRD specific binding regions. Fibril formation and oligomerization can have a major impact on the availability of the PRD for protein-protein interactions (Ko et al., 2018; Posey et al., 2018). It has previously been proposed that the high density of surface-exposed bristles (such as those in P and T-fibrils) can cause increased binding affinities for PRD binding proteins such as profilin (Posey et al., 2018). We made similar observations with antibodies that bind to the PRD, which had a much stronger binding affinity to unbundled fibrils than to monomeric HTTex1 (Ko et al., 2018). Thus, fibrils or oligomers could further contribute to toxicity by having enhanced binding affinities to binding partners, which could cause mislocalization of HTTex1 binding partners. This notion is consistent with the fact that several PRD binding proteins have been shown to colocalize with cellular huntingtin aggregates (Qin, 2004; Ralser et al., 2005; Shirasaki et al., 2012; Sittler et al., 1998). Increased binding affinities to T-fibrils, P-fibrils or oligomers could cause mislocalization and misregulation of such proteins. In contrast, the N-fibrils with their more tightly packed PRD regions are less likely to interact with such binding partners. In fact, we recently reported that PRD binding antibodies have a very weak affinity for the N-fibrils (Ko et al., 2018).

From a methodological perspective, the combined use of EPR distances, EPR mobilities and solid-state NMR may well allow us to obtain a detailed three-dimensional structural model of the different fibril types, including detailed local structure for the N17 and polyQ region region. Such structural information should prove useful for the design molecules aimed at shifting the structural states of toxic species into nontoxic ones.

## Online Methods

### Protein expression and purification

HTTex1(Q46) was recombinantly expressed as a thioredoxin fusion (Trx-HTTex1) using a pET32a vector in BL21 (DE3) *E.coli.* The expression and purification of uniformly ^15^N-^13^C labeled HTTex1(Q46) for NMR measurements and non-isotope labeled Cys mutants for EPR measurements was performed using His60 resin (Clontech) and a subsequent HiTrap Q XL anion exchange chromatography column (GE Healthcare) on an AKTA FPLC system (Amersham Pharmacia Biotech) as previously described (Bugg et al., 2012; Isas et al., 2017).

### Fibril formation and seed prepration

HTTex1 T-fibrils for NMR and EPR measurements were made by starting with the Trx-HTTex1 fusion proteins at a concentration of 25 μM (632 μg/ml) in 20 mM Tris, 150 mM NaCl with 5% molar ratio of HTTex1(Q46) seeds added. Fibril started forming after adding 1 unit of EKMax per 280 μg of protein. The reaction mixture was left without agitation at 4°C for 1 weeks to complete fibril formation. HTTex1 N-fibrils were prepared similarly but by incubating them at 37°C instead of 4°C.

N-fibrils were transformed into T-fibrils by pelleting the fibrils (60 min at 150,000g) and redissolving the pellet in 0.5% TFA in H_2_O. The fibrils could be further disaggregated by adding an additional washing step with HFIP. Here the N-fibril pellet was redissolved in HFIP. The HFIP was consequently evaporated under a stream of N2 and the dried fibril pellet was again redissolved in 0.5% TFA. This treatment resulted in what we term P-fibrils.

HTTex1 seeds were prepared as described previously.(Isas et al., 2017) In short, seed formation was initiated by removing the Trx fusion tag using 1 unit EKMax (Invitrogen) per 280 μg of HTTex1. The mixture was kept at 4°C for several days and sonicated. Seeds were analyzed using electron microscopy for morphology and a BCA assay (Thermo Fisher Scientific) was used to determine their concentration. This procedure was also used to produce sonicated fibrils for dot bot analysis.

### Electron microscopy

Grids for electron microscopy (150 mesh copper) were prepared by pipetting 10 μl of sample on them and letting the sample absorb for 5 to 10 minutes. For negative stain, 10 μl of 1% uranyl acetate solution was added and the grids were incubated for 2 min and then dried at room temperature for 1 hour. All grids were captured on a Gatan digital camera using a JEOL JEM-1400 electron microscope (JEOL, Peabody, MA) at 100 kV.

### Circular Dichroism

CD spectra of HTTex1 fibrils were measured at a concentration of 5-20 μM in 10 mM phosphate buffer with no salt, pH 7.4. All CD spectra were acquired on a Jasco 815 spectropolarimeter (Jasco, Inc., Easton, MD). Data points were measured every 0.5 nm at a scan speed of 50 nm/min from 260 to 190 nm. Between 20–30 acquisitions were averaged for each sample. Background spectra of buffer only were subtracted to obtain the final spectrum. Temperature dependence of the CD spectra was measured using a Jasco PFD-425s temperature controller. PolyP solutions were incubated at a given target temperature for at least 20 min before recording the CD spectra. All experimental details stayed as described above. The difference spectra of N and T-fibrils and polyP CD spectra recorded at 90 and 25°C were smoothed using the Savitzky-Golay method in scientific python.

### EPR spectroscopy

HTTex1 fibrils were sedimented using ultracentrifugation, resuspended in 20 μl of 20 mM Tris, 150 mM NaCl, and filled into quartz capillaries (ID=0.6 mm and OD=0.84 mm, VitroCom, Mt. Lakes, NJ). EPR spectra were recorded on an X-band Bruker EMX spectrometer (Bruker Biospin) at ambient temperature. The sweep width was 150 G using a HS cavity at an incident microwave power of 12.60 mW. All spectra were normalized by double integration.

Kinetic studies of fibril formation were done as described previously (Pandey et al., 2018). In short, a 15 μM solution of HTTex1 spin labeled at residue 35 (35R1) was mixed with HTTex1(Q46) P, T, or N-fibril seeds at a 5% molar ratio or no seeds. The sample was then loaded into quartz capillaries and a EPR spectrum was recorded every 15 minute for 17.5 hours using the parameters described above.

To determine the distance between spin labels, four pulse DEER experiments (Pannier et al., 2000) were measured at a temperature of 78 K on a Bruker Elexsys E580 X-band pulse EPR spectrometer equipped with a 3 mm split ring (MS-3) resonator, a continuous-flow cryostat (CF935, Oxford Instruments), and a temperature controller (ITC503S, Oxford Instruments). Trx-HTTex1 fusion protein with spin label pairs were diluted with 90-95% non-spin labeled Trx-HTTex1 before inducing fibril formation with EKMax to reduce background from intermolecular distances. For some samples 15-20% sucrose was added as cryoprotectant after fibril formation. The fibril samples were flash frozen and measured. The data were fitted using a single Gaussian as implemented in DEER Analysis 2019 (Jeschke et al., 2006).

### Solid-state NMR

All solid-state NMR experiments were acquired on a 600 MHz Agilent DD2 spectrometer using a 1.6 triple-resonance probe operating at 25 kHz MAS, 0°C. Hard pulses nutation frequencies for ^1^H and ^13^C were 200 kHz and 100 kHz respectively. ^1^H-^13^C Hartmann-Hahn cross polarization was done with using spin-lock nutation frequencies of 85 kHz for ^1^H and 60 kHz for ^13^C with a 10% ramp in ^1^H radio-frequency (rf) amplitude. Two pulse phase modulation (TPPM) ^1^H decoupling with an RF field strength of 120 kHz was used during indirect and direct acquisitions. A pre acquisition delay of 3 s was used for all NMR experiments. One dimensional ^13^C CP and refocused INEPT (insensitive nuclei enhanced by polarization transfer) spectra were recorded with a spectral width of 50 kHz and 650 complex points. For the fibrils prepared at 4°C and 37°C 256 scans were acquired. The 1D spectra of the protofibrils were acquired with 1024 scans. 2D DARR (dipolar assisted rotational resonance)(Takegoshi et al., 2001) spectra were recorded with a 25 kHz ^1^H spin-lock recoupling rf-field, a mixing time of 50 ms, and 50 kHz spectral widths in both dimensions with 650 complex points in the direct and 300 complex points in the indirect dimension. The DARR spectra of fibrils prepared a 4°C, 37°C, protofibril, were recorded with 16, 32, and 48 scans for each indirect increments, respectively. ^1^H-^13^C INEPT heteronuclear correlation (HETCOR) spectra were recorded with a ^13^C refocusing pulse in the indirect ^1^H dimension, which had a spectral width of 10 kHz, and was sampled with 16 scans for each of the 128 complex points. The direct ^13^C dimension had a spectral width of 50 kHz and was sampled with 650 complex points. All spectra were referenced to 4,4-dimethyl-4-silapentane-1-sulfonic acid (DSS) using adamantane as an external standard.(Harris et al., 2008)

### Dot Blot

HTTex1 fibril and monomer samples (1.5 μl per sample) were blotted on a nitrocelluose membrane and incubated with the desired primary antibody at a 1:5000 dilution. The MW8 antibody was a gift from Ali Khoshnan, the monoclonal Anti-polyHisitinde antibody was purchase from Sigma-Aldrich. A fluorescently-labeled secondary antibody (IRDye-800 anit-Mouse IgG) was used at a dilution of 1:10,000. The dot blot was visualize using a LI-COR Odyssey Infrared Imaging system.

### Cellular Puncta Formation Assay

Neuro2a cells, grown on glass dishes coated with Matrigel, were transfected with plasmid constructs using a Lipofectamine LTX with Plus Reagent (ThermoFisher) transfection kit according to manufacturer protocol. The plasmid constructs containing HTTex1 with C-terminal EGFP and polyQ repeat lengths of Q16, Q39, Q72 or EGFP alone as control were described previously (Fodale et al., 2014). Co-transfection was performed in the same manner described by Nekooki-Machida et al. with the addition of 2.5 μg of fibrils 24 hours after plating using Lipofectamine LTX. BSA fibrils were used as a control. To make BSA fibrils, lyophilized BSA was dissolved in PBS at a concentration of 100 μM, incubated for 1 h at 40°C, sonicated, and stored in 4°C until use. To confirm the intracellular delivery of protein in the co-transfection protocol, HTTex1 labeled with Alexa Fluor 594 at residue 101 was transfected and its location confirmed by fluorescence microscopy. Neuro2a culture were incubated for 48 hours and imaged live using a Zeiss AxioPlan2 microscope. Fluorescent and differential interference contrast (DIC) images were acquired using a 40x magnification. For each condition the experiment was repeated three times and images of 3 visual fields were taken and analyzed for each repeat. The total number of transfected cells expressing EGFP and number of cells with EGFP foci of huntingtin aggregates was counted in a blinded fashion. Two-sided student T-tests were used for statistical analysis to compare the various conditions vs control.

