## Supplemental Figures S1-S4 for "Huntingtin fibrils with different toxicity, structure, and seeding potential can be reversibly interconverted"

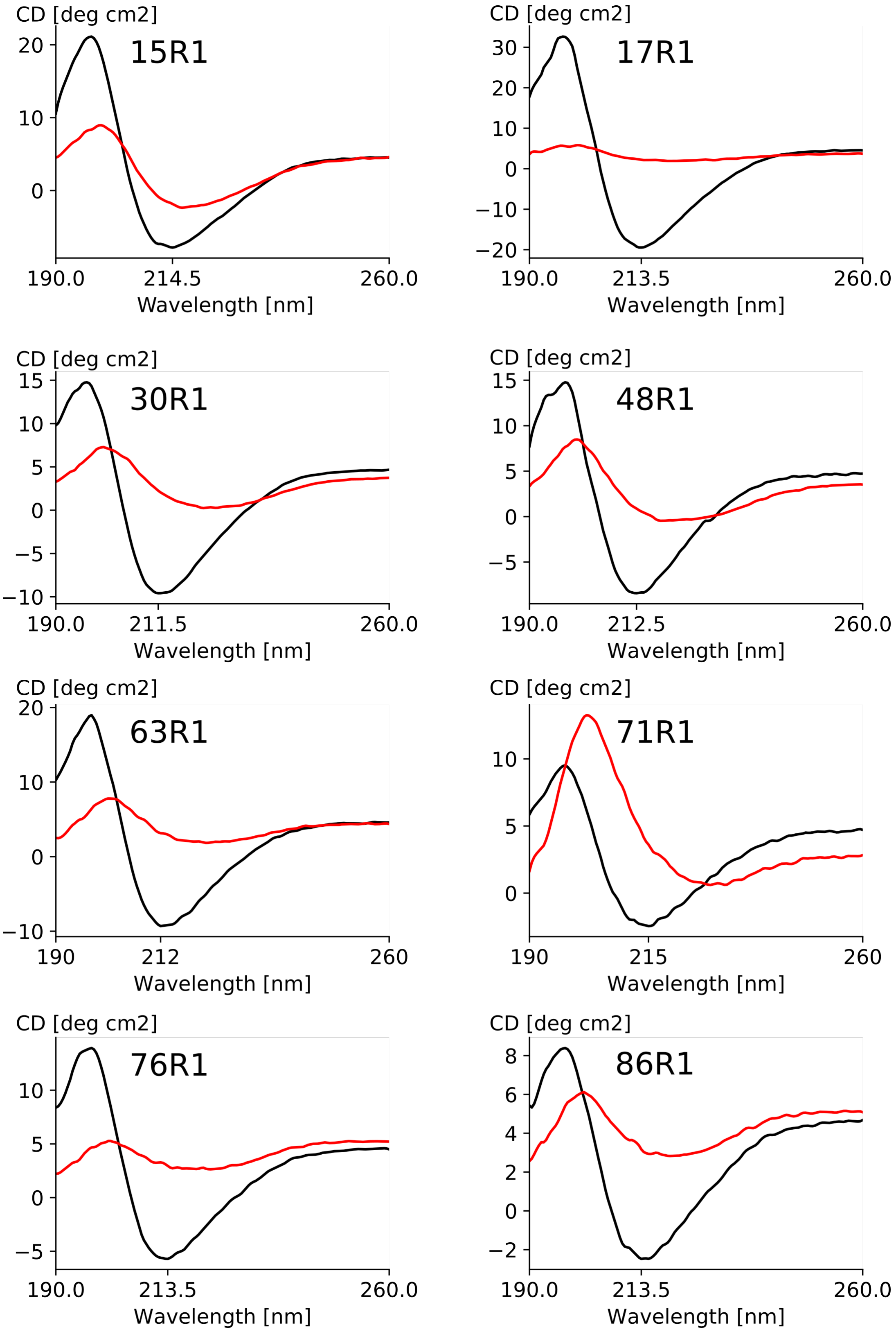


**Supplementary Figure S1:** CD spectra of all the MTSL labeled HTTex1 samples used for the EPR data of Figure 2. The T-fibril spectra (black) show typical minima around 213 nm whereas all the N-fibril spectra (red) are red shifted and are of decreased intensity because the bundling of the fibrils increases scattering.

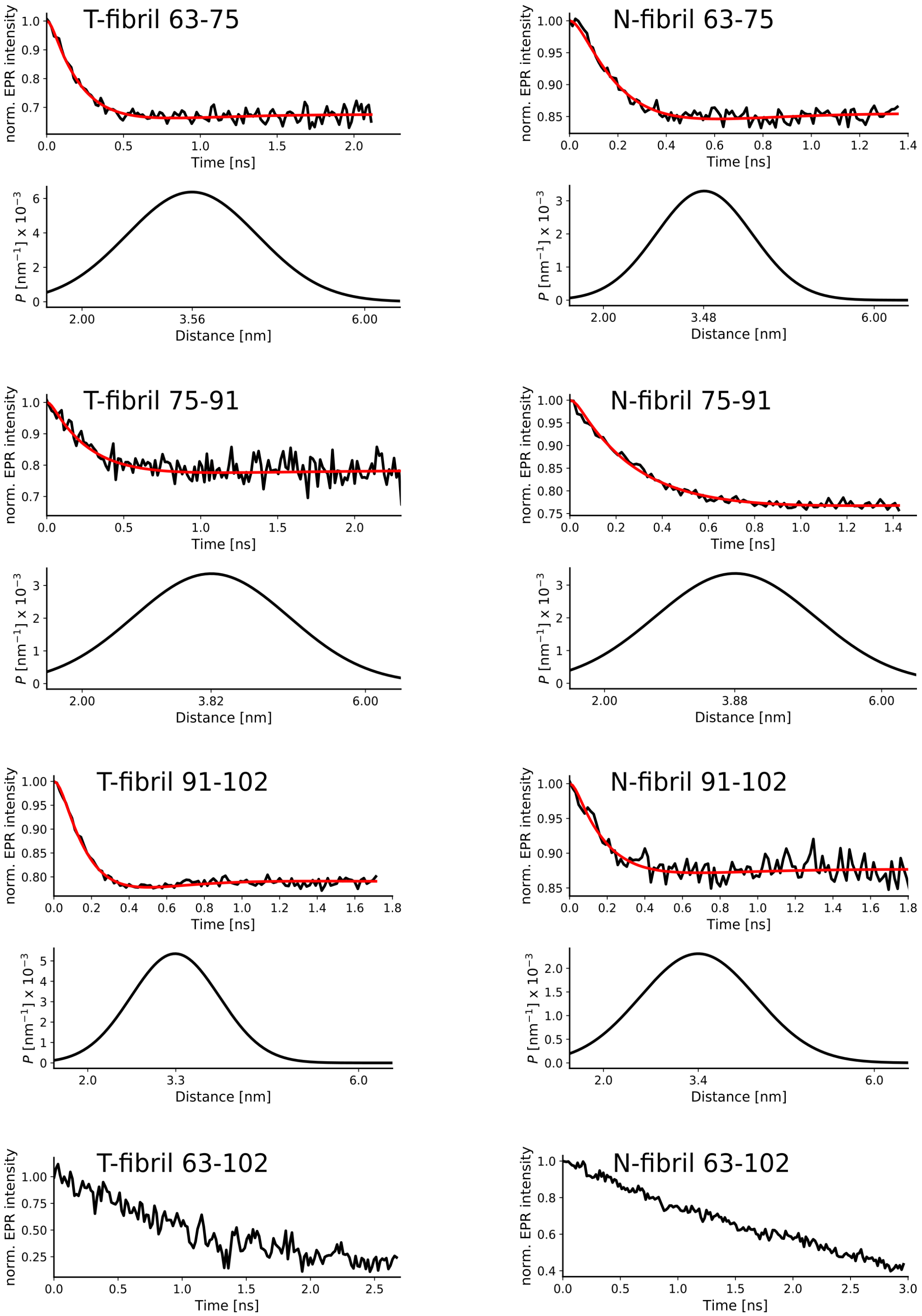


**Supplementary Figure S2:** DEER data and fits corresponding to the distances reported in Figure 5. Top panels: base line corrected DEER data (black) and fit to single Gaussian distribution (red). Bottom panels: Gaussian distribution corresponding to fit. The center of the distribution is indicated. Because the distance between residues 63-102 was above the detection limit, no fit is shown and no baseline was substracted in this case.

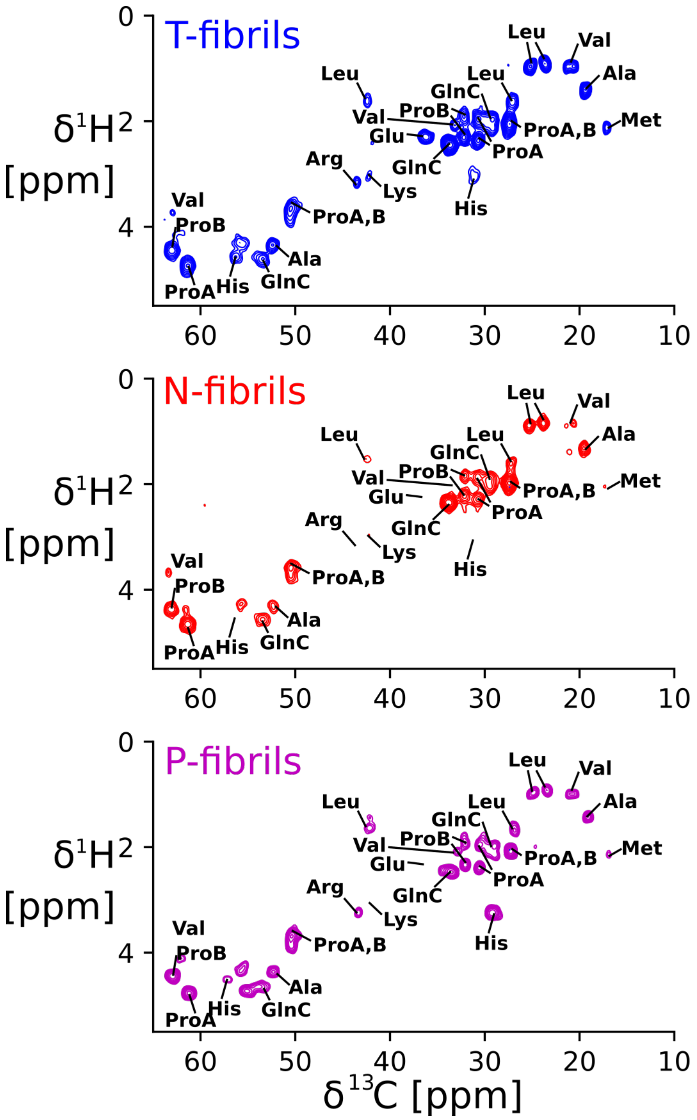


**Supplementary Figure S3:** INEPT HETCOR 2D $^{1}$H-$^{13}$C spectra of the three HTT$_{ex1}$Q46 fibril types show very little chemical shift changes. Notable differences include the absence of His shifts in the N-fibrils and the shift of the corresponding peaks in the protofibril spectra due to a change in pH. While the extent of dynamics in the C-termini varies between the different fibril types, their chemical shift does not change.

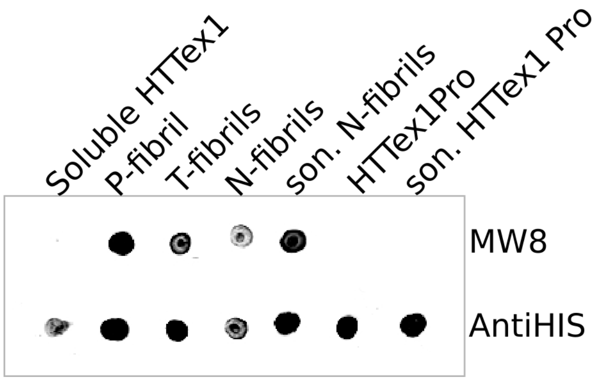


**Supplementary Figure S4:** The exposed PRD of the T and P-fibrils is better accessible PRD specific antibodies than the PRD of the N-fibrils. Dot blot of HTTex1 monomer and different HTTex1 fibril types. The MW8 and fibril specific antibody that recognizes an epitope in the PRD and an His tag antibody were used for detection. MW8 strongly binds to P and T-fibrils and to a lesser degree N-fibrils. Sonication of N-fibrils (son. N-fibril) significantly increases the binding of MW8. The HTTex1 monomer and fibrils formed by a HTTex1 mutant lacking the MW8 epitope of the PRD (HTTex1Pro) show no binding to MW8.
